# Common as well as unique methylation-sensitive DNA regulatory elements in three mammalian SLC9C1 genes

**DOI:** 10.1101/2023.08.29.555319

**Authors:** Cameron C. Gardner, Jason A. Abele, Thomas J. Winkler, Caroline N. Reckers, Sydney A. Anas, Paul F. James

## Abstract

The SLC9C1 gene (which encodes the NHE10 protein) is essential for male fertility in both mice and humans, however the epigenetic mechanisms regulating its testis/sperm-specific gene expression have yet to be studied. Here we identify and characterize DNA regulatory elements of the SLC9C1 gene across three mammalian species: mouse, rat, and human. First, *in silico* analysis of these mammalian SLC9C1 genes identified a CpG island located upstream of the transcription start site in the same relative position in all three genes. Further analysis reveals that this CpG island behaves differently, with respect to gene regulatory activity, in the mouse SLC9C1 gene than it does in the rat and human SLC9C1 gene. The mouse SLC9C1 CpG island displays strong promoter activity by itself and seems to have a stronger gene regulatory effect than either the rat or human SLC9C1 CpG islands. While the function of the upstream SLC9C1 CpG island may be divergent across the three studied species, it appears that the promoters of these three mammalian SLC9C1 genes share similar DNA methylation-sensitive regulatory mechanisms. All three SLC9C1 promoter regions are differentially methylated in lung and testis, being more hypermethylated in lung relative to the testis, and DNA sequence alignments provide strong evidence of primary sequence conservation. Luciferase assays reveal that *in vitro* methylation of constructs containing different elements of the three SLC9C1 genes largely exhibit methylation-sensitive promoter activity (reduced promoter activity when methylated) in both HEK 293 and GC-1spg cells. In total, our data suggest that the DNA methylation-sensitive elements of the mouse, rat, and human SLC9C1 promoters are largely conserved, while the upstream SLC9C1 CpG island common to all three species seems to perform a different function in mouse than it does in rat and human. This work provides evidence that while homologous genes can all be regulated by DNA methylation-dependent epigenetic mechanisms, the location of the specific cis-regulatory elements responsible for this regulation can differ across species.

## Introduction

The Na^+^/H^+^ exchangers (NHEs) represent a family of ion transporter proteins that function to regulate pH and osmolarity in various cellular compartments across many different tissues and cell types. Most characterized NHE proteins localize to the plasma membrane where they function to regulate intracellular pH (pH_i_) by exporting intracellular protons (H^+^) using the energy stored in the inward directed sodium ion (Na^+^) electrochemical gradient established by the Na,K-ATPase (Orlowski and Grinstein, 2004). Intracellular pH has long been known to be important for a wide range of physiological processes including sperm motility (Babcock et al., 1983; Lee, 1984; Nishigaki et al., 2014) as pH likely affects multiple proteins vital for sperm function such as the Ca^2+^ channel complex CatSper and the K^+^ channel SLO3 (Brenker et al., 2014; Brukman et al., 2019; Chávez et al., 2014; Hwang et al., 2019; Kang et al., 2021; Kirichok et al., 2006). Studies have shown that the transition from the immotile to the motile state and/or from a less motile to a highly motile state of rat and bovine sperm is mediated through a rise in pH_i_ (Babcock et al., 1983; Carr and Acott, 1989; Matamoros-Volante and Trevino, 2020; Vijayaraghavan et al., 1985; Wong et al., 1981). In fact, sperm pH_i_ positively correlates with both hyperactivated motility and successful *in vitro* fertilization (IVF) in normospermic human patients (Gunderson et al., 2021).

Na^+^/H^+^ Exchangers are encoded by the SLC9 gene family of solute carriers. This gene family has been divided into three subfamilies, determined by sequence homology: the NHE subfamily (SLC9A1-SLC9A9 which encode NHE1- NHE9), the NHA subfamily (SLC9B1 and SLC9B2 which encode NHA1 and NHA2 [also known as NHEDC1 and NHEDC2]), and the mammalian sperm-NHE-like subfamily (SLC9C1 and SLC9C2 which encode NHE10 [also known as sNHE] and the NHE11 protein) (Donowitz et al., 2013; Gardner and James, 2023; Pedersen and Counillon, 2019). The SLC9C1 gene (encoding the NHE10 protein) was first characterized in mice to be a testis/sperm-specific expressed gene whose protein product localizes to the sperm principal piece (Wang et al., 2003). Among the SLC9C proteins, NHE10 and NHE11 share a unique predicted protein structure consisting of an NHE domain at the N-terminus, followed by a voltage-sensing domain (VSD), and an intracellular cyclic nucleotide binding domain (CNBD) at the C-terminus (Brett et al., 2005; Gardner and James, 2023; Pedersen and Counillon, 2019; Wang et al., 2007, 2003). The transport activity of the mammalian SLC9C proteins (NHE10 and NHE11) have yet to be characterized. However, the singular sea urchin SLC9C protein (originally suggested to be the NHE10 isoform) which also possesses a similar structure to mammalian NHE10 and NHE11 of having a NHE domain, VSD, and CNBD was shown to exchange Na^+^ for H^+^ thus regulating pH_i_, while also being finely regulated by both membrane potential (E_m_) and cyclic nucleotides (Windler et al., 2018). The ability for an SLC9C protein to be regulated by changes in membrane potential is important in the context of mammalian sperm physiology because sperm hyperpolarize their E_m_ following capacitation via a pH-mediated K^+^ efflux through the SLO3 channel (Balbach et al., 2020; Brenker et al., 2014; Brown et al., 2016; Chávez et al., 2014; López-González et al., 2014; Santi et al., 2010). Additionally, cAMP (primarily synthesized by soluble adenylyl cyclase (sAC)) is known to be important for the initiation of sperm motility upon release from the epididymis, induction of hyperactivated motility, and increased protein tyrosine phosphorylation allowing sperm to undergo acrosomal exocytosis and for rendering the spermatozoa competent for fertilization (Balbach et al., 2020; Jansen et al., 2015; Marquez and Suarez, 2004; Naz and Rajesh, 2004; Visconti et al., 1995; Zippin et al., 2001). Thus, regulation by these sperm physiological signals allows for fine-tuning of NHE10 and/or NHE11 activity to meet specific environmental cues of the sperm in its journey towards fertilization. NHE10 is thought to be the protein involved in regulating the hyperpolarization-mediated cytosolic alkalization that is observed in the flagellum of mouse sperm through its voltage sensing domain (Hernandez- Garduno et al., 2022), highlighting NHE10’s importance in the mouse sperm capacitation process.

Mice lacking the NHE10 protein (encoded by SLC9C1) are completely infertile due to immotile sperm (Wang et al., 2003) and it was found that NHE10 protein expression is necessary for full-length soluble adenylyl cyclase protein expression (Wang et al., 2007), which is another protein essential for male fertility in both mice and humans (Akbari et al., 2019; Esposito et al., 2004; Hess et al., 2005). A clinical study of subfertile human male patients found significantly less NHE10 protein in sperm from asthenozoospermic compared to normozoospermic men and that NHE10 expression was positively correlated with higher sperm motility parameters (Zhang et al., 2017). A recent analysis of an infertile male presenting with asthenozoospermia found that this patient bares a homozygous mutation in the SLC9C1 gene predicted to generate a truncated NHE10 protein (Cavarocchi et al., 2021). SLC9C1 was determined to be expressed exclusively in the testis in mice via a full tissue RNA dot-blot assay (Wang et al., 2003) and RNA sequencing suggests that SLC9C1 gene expression is also limited to the testis in both rat https://www.ncbi.nlm.nih.gov/gene/288117/ and human https://www.ncbi.nlm.nih.gov/gene/285335. Although the SLC9C1 gene is clearly important for male fertility in both mice and humans, the epigenetic mechanisms by which the SLC9C1 gene regulates its testis/sperm-specific expression has not yet been studied. Previous work in our lab has demonstrated that both the human SLC9B1 gene and the mouse ATP1A4 gene, contain regulatory elements that may help ensure their testis/sperm-limited expression through DNA methylation-dependent epigenetic mechanisms (Kumar et al., 2016; Kumar and James, 2015). Here we examine whether the testis/sperm-specific expression of SLC9C1 can also be influenced by DNA methylation.

DNA methylation in mammals is an epigenetic mechanism involving the addition of a methyl group to the cytosine in a CpG dinucleotide context by enzymes called DNA methyltransferases (DNMTs) (Suzuki and Bird, 2008). Located sparsely throughout vertebrate genomes are CpG-rich regions called CpG islands (CpGI). These CpG islands are characterized as being on average around 1000 base pairs (bp) in length, enriched in G+C base composition, and are often lacking DNA methylation (Deaton and Bird, 2011). CpG islands can be characterized as promoter CpG islands, intragenic CpG islands (found within a gene body), or intergenic CpG islands (found between gene bodies). Promoter CpG islands have been studied extensively and have largely been characterized as substantial epigenetic regulators of gene expression; DNA methylation alters transcription factor binding to promoter regions, affecting RNA polymerase binding and subsequent transcription. Both intragenic and intergenic CpG islands have been shown to sometimes act as alternative promoters themselves (Deaton and Bird, 2011). Although methylated DNA has canonically been associated with repressed gene expression, more recent insight has found that methylated DNA can also be associated with increased gene expression, especially when intragenic CpG islands are methylated (Greenberg and Bourc’his, 2019; Spruijt and Vermeulen, 2014). Clearly DNA methylation plays an important epigenetic role, however whether DNA methylation increases or decreases gene expression appears to be dependent on the specific cellular and chromatin context (Wan et al., 2015).

DNA methylation has been shown to be an important epigenetic regulator of various tissue-specific expressed genes, including those in the germline (Allegrucci et al., 2005; Ben Maamar et al., 2022). Some examples are the rat testis-specific H2B (TH2B) histone gene and the mouse ALF gene, which both express germ-cell specific transcripts. Both promoters are methylated in somatic tissue but are unmethylated in the male germ cells where these genes are expressed (Choi and Chae, 1991; Xie et al., 2002). Another example is the ZNF645 gene whose expression was shown to be regulated by demethylation of the promoter region in testis and in some lung cancerous tissues while the promoter remained methylated in healthy lung tissues where it is not expressed (Bai et al., 2010). Furthermore, it has been shown that methylation of a specific CpG dinucleotide located within an activating transcription factor/cAMP-responsive element-binding (ATF/CREB) site located just upstream of the Pdha-2 gene promoter is responsible for inhibiting transcription in non-male germ cells (Iannello et al., 2000). DNA methylation of the CpG island overlapping the promoter region of the human SLC9B1 gene has been shown to strongly regulate gene expression and contributes to this gene’s testis/sperm- restricted expression (Kumar and James, 2015). The mouse ATP1A4 gene has also been demonstrated to be regulated by DNA methylation; both the promoter and an intragenic CpG island are methylation-sensitive and likely contribute to its testis/sperm-specific expression (Kumar et al., 2016). DNA methylation is critically important for male fertility because it was recently shown that DNMT3A expression, an isoform of one of the DNA methylation enzymes, is required for spermatogonial stem cells (SSCs) to commit to spermatogenesis and differentiate (Dura et al., 2022). Finally, numerous studies have shown that aberrant DNA methylation of the promoter sequences of human testis-expressed genes in male patients are associated with infertility (Gunes et al., 2016; Han et al., 2020; Hung et al., 2021; Richardson et al., 2014; Sugimoto et al., 2009; Tang et al., 2018; Xu et al., 2016).

In this study we compare the role that DNA methylation plays in regulating the expression of the SLC9C1 gene in three mammalian species: mouse, rat, and human. *In silico* analysis revealed that although all possess intragenic CpG islands, none of these intragenic CpG islands shared a common location within the gene in all three species. However, there is a conserved CpG island located ∼1000 bp upstream of the transcription start site of the SLC9C1 gene in each species. This upstream CpG island as well as the putative promoter regions of the three SLC9C1 genes were analyzed by bisulfite sequencing and *in vitro* luciferase reporter assays to determine if these DNA elements could contribute to the tissue-specific expression of this gene via DNA methylation. Bisulfite sequencing analysis revealed that both the mouse SLC9C1 promoter and CpG island were very clearly differentially methylated between the lung (a non-expressing tissue) and the testis (an expressing tissue). The rat and human SLC9C1 promoters showed less differential methylation between the testis and the lung than the mouse promoter did, however, both the rat and human promoters displayed increased methylation in the lung compared to the testis. Unlike the mouse, there was no difference in the methylation of the rat and human CpG islands between lung and testis. Luciferase reporter assays revealed differences in the strength and methylation-sensitivity of the three species’ promoters in somatic and spermatogonial cells in culture. These reporter assays also suggest that the mouse SLC9C1 CpG island plays a different role in regulating SLC9C1 gene expression than do the rat and human SLC9C1 CpG islands.

## Methods

### *In silico* Genomic DNA analyses

The SLC9C1 promoter sequences were determined using the eukaryotic promoter database (Dreos et al., 2017) and cross-referenced with DeePromoter (Oubounyt et al., 2019) and potential promoters that were recognized by both programs were used in our analyses. The entire NCBI annotated SLC9C1 genes (*Mus musculus* SLC9C1 Gene ID: 208169, *Rattus norvegicus* SLC9C1 Gene ID: 288117, *Homo sapiens* SLC9C1 Gene ID: 285335) including 10,000 bp upstream of the predicted transcription start site of each gene were analyzed for potential CpG islands using the DBCAT CpG island Finder software (http://dbcat.cgm.ntu.edu.tw/) and the GenScript Sequence Manipulation Suite CpG Islands software (Stothard, 2000) with the default settings for both programs.

### DNA methylation analysis using bisulfite sequencing

Genomic DNA was extracted from mouse (*Mus musculus*) or rat (*Rattus norvegicus*) lung and testis tissue by phenol-chloroform extraction followed by ethanol precipitation. Human lung and testis genomic DNA from normal adult tissue was purchased (BioChain). The EZ DNA Methylation-Gold Kit (Zymo Research) was used to treat 1000 ng of genomic DNA according to manufacturer instructions. The primers that were used to amplify the bisulfite treated genomic DNA were designed using the MethPrimer software (Li and Dahiya, 2002) (Table 1). The DNA fragments were PCR amplified with the Q5U DNA polymerase (New England BioLabs) and then gel purified using the Gel/PCR DNA Fragments Extraction Kit (IBI Scientific). The gel purified DNA fragments were cloned into the pMiniT 2.0 vector using the NEB PCR Cloning Kit (New England BioLabs) and then transformed into NEB 10-beta competent cells (New England BioLabs). Plasmid DNA was extracted from ten positive clones using the GeneJET Plasmid Miniprep Kit (Thermo Scientific) and Sanger sequenced (EuroFins Genomics). The methylation status of these individual CpG clones were determined with the Quantification Tool for Methylation Analysis (QUMA) software (Kumaki et al., 2008).

**Table 1.**
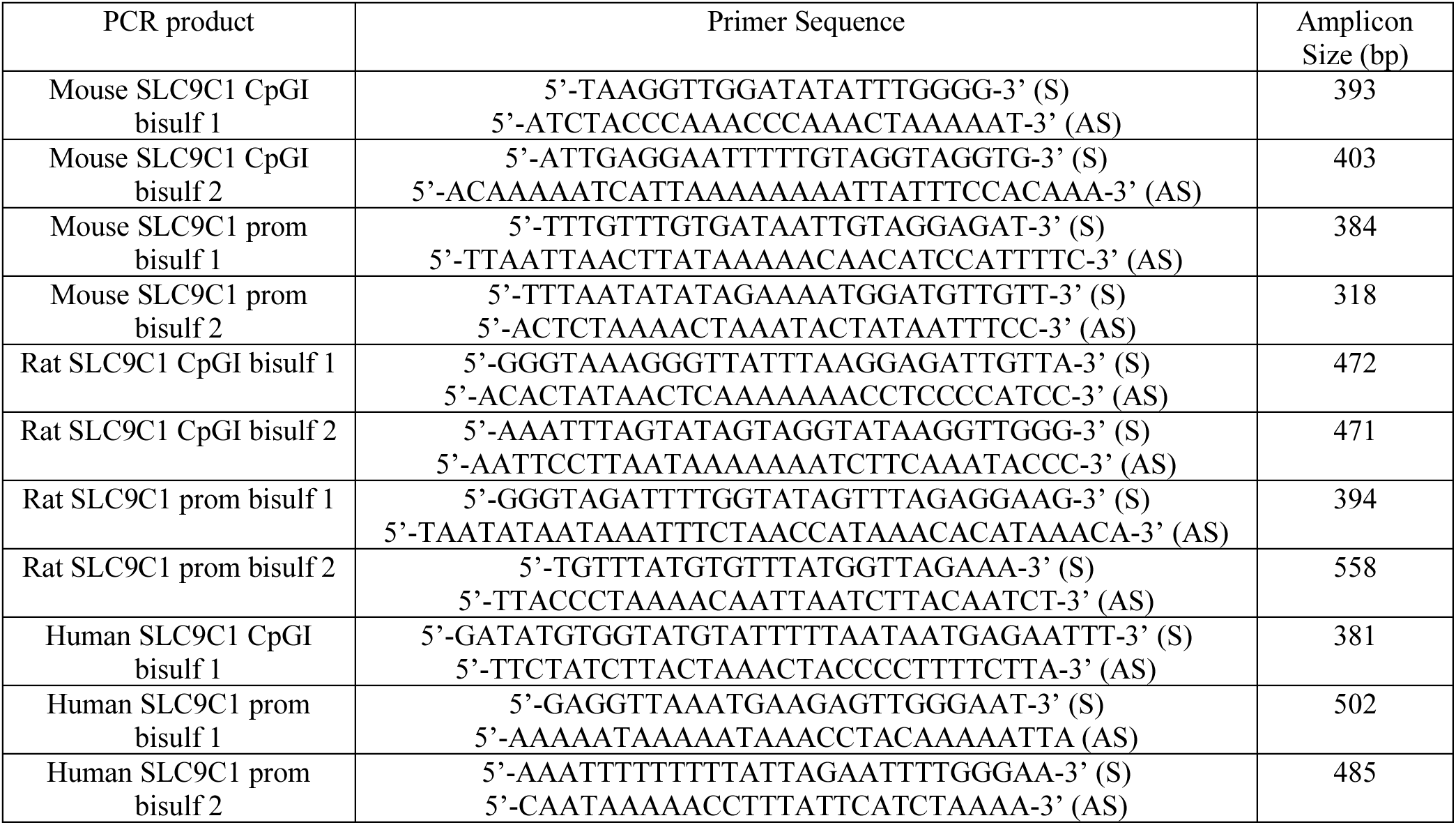
Primers used in this study to amplify bisulfite treated genomic DNA for methylation analysis.

### Generation of pCpGL luciferase reporter constructs

The pCpGL-Basic vector (a kind gift of the Rehli lab) was used as the vector backbone for all tested firefly luciferase reporter constructs. The pCpGL-Basic vector does not possess any CpG dinucleotides and therefore is useful for studying the effect of DNA methylation on promoter and enhancer/silencer activity (Klug and Rehli, 2006). The promoters and CpG islands of the three studied species were first PCR amplified with Q5 DNA polymerase (New England BioLabs) using gene-specific primers (Table 2) from lung genomic DNA and then gel purified using the Gel/PCR DNA Fragments Extraction Kit (IBI Scientific). The gel purified DNA fragments were cloned into the pMiniT 2.0 vector, transformed into NEB 10-beta competent cells, and plasmid DNA was extracted and sequenced as described above. Then the regions of interest of the CpG island and promoters were amplified using Q5 DNA polymerase with primers which possess complementary overhangs to the pCpGL-Basic vector (complementary to the multiple cloning site just upstream of the firefly luciferase open reading frame) (Table 3). These PCR amplicons were gel purified and then assembled using NEBuilder HiFi DNA Assembly Master Mix (New England BioLabs) and transformed into ChemiComp GT115 competent cells (InvivoGen). The correct insertion of all constructs was verified by Sanger sequencing. The pCpGL-CMV (containing the minimal CMV promoter) and pCpGL-EF1α (containing the human EF1α promoter plus its first intron) (Kumar et al., 2016; Kumar and James, 2015) were used to generate reporter constructs with the CpG islands of the SLC9C1 genes upstream of the CMV and EF1α promoters.

**Table 2.**
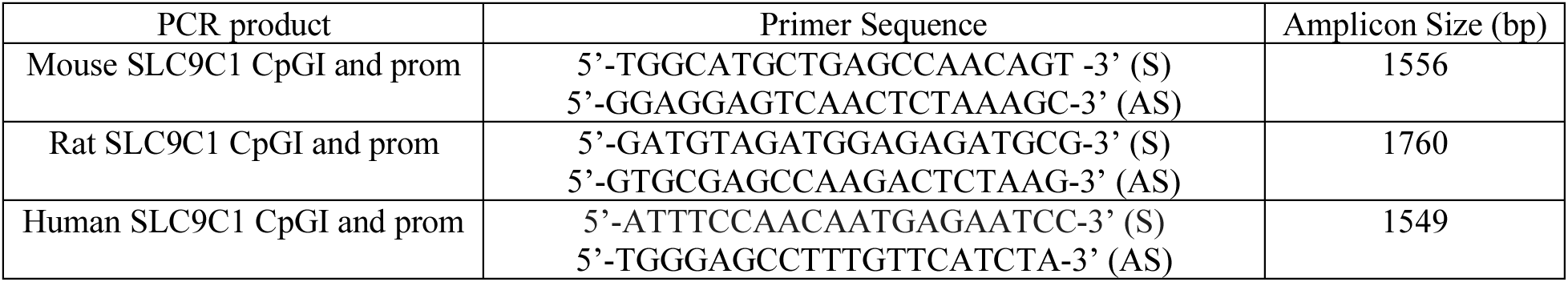
Primers used in this study to amplify different genomic regions of the mouse, rat, and human SLC9C1 gene. These PCR products were first PCR cloned into pMiniT2.0 to verify sequences.

**Table 3.**
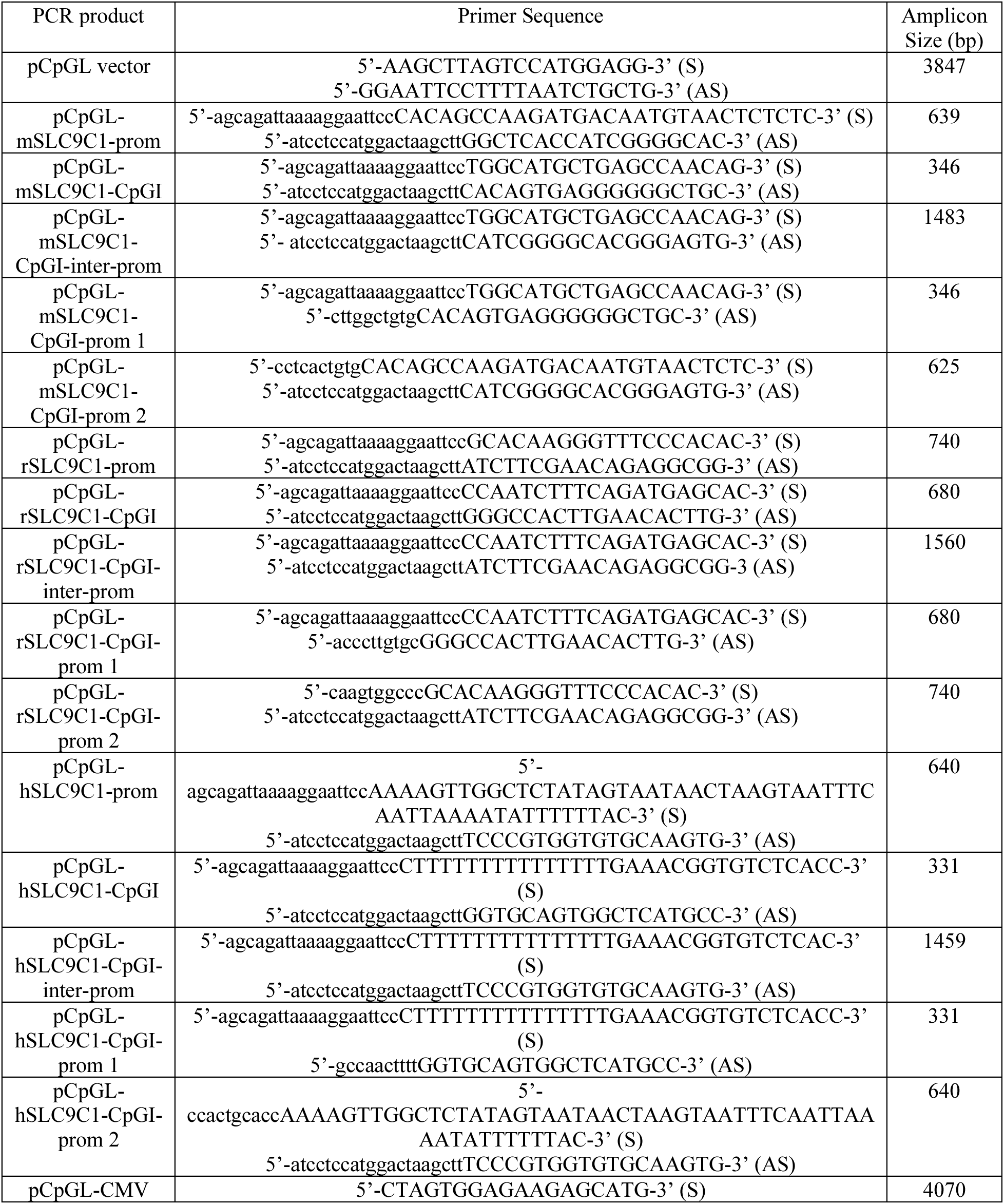
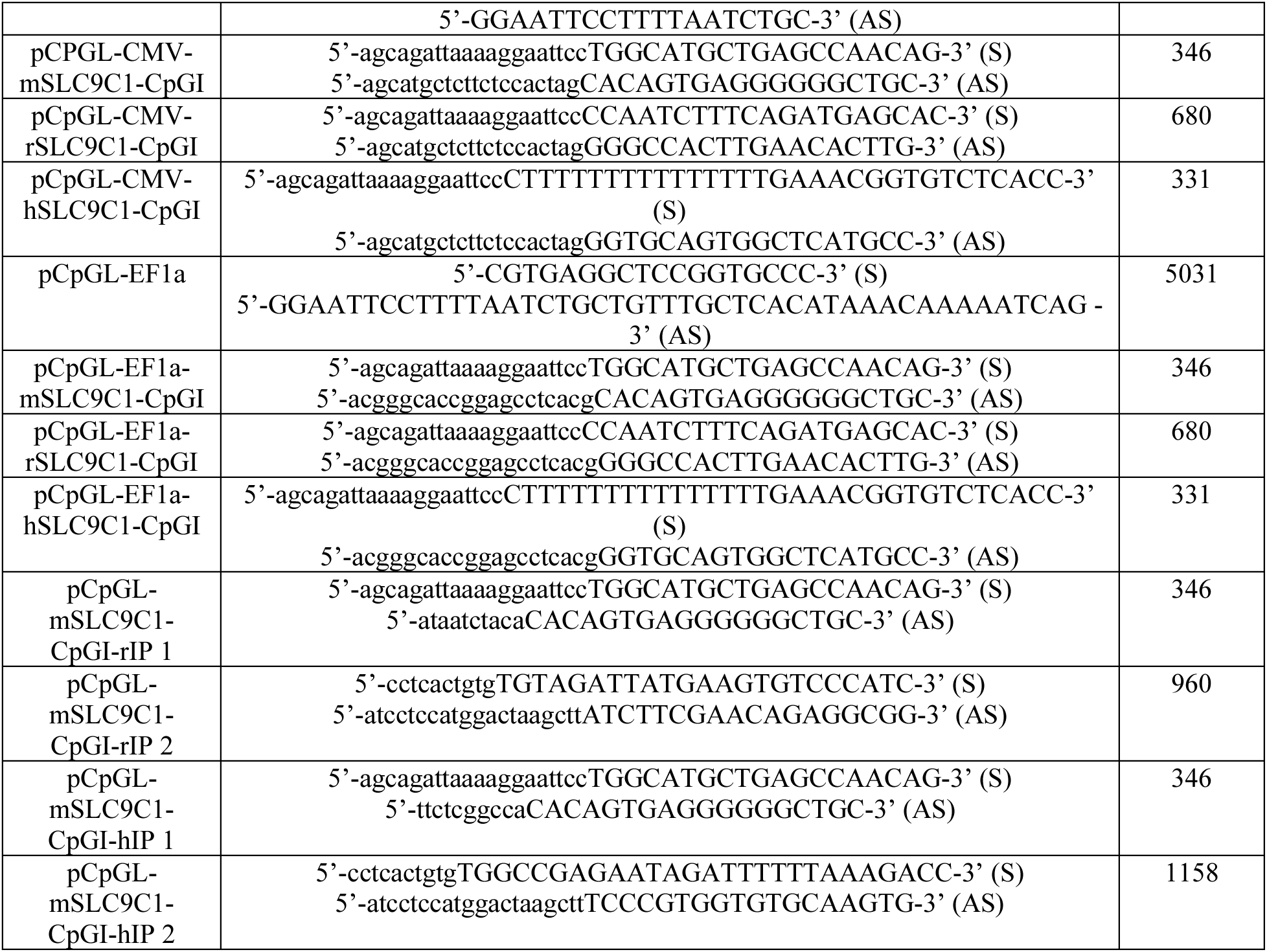
Primers used in this study to amplify and assemble different genomic regions of the mouse, rat, and human SLC9C1 gene into the pCpGL vector. All inserts were cloned into the pCpGL vector using the HiFi DNA Assembly. The uppercase letters in the primer sequences indicate gene specific sequences and the lowercase letters indicate sequences complementary to the target vector into which they were assembled.

### *In vitro* methylation of plasmid DNA

Plasmids were *in vitro* methylated using the M. SssI CpG methyltransferase enzyme (New England BioLabs) according to manufacturer recommendations. In brief, each construct was incubated with 10 units of M. SssI enzyme in the presence of 160 µM of the methyl donor S-adenosylmethionine (New England BioLabs) for 4 hours at 37°C. The mock methylated constructs were incubated under the same conditions but in the absence of the M.SssI methyltransferase enzyme. All reactions were heat inactivated at 65°C for 20 min and then the DNA was purified by ethanol precipitation. The ethanol precipitated DNA constructs were digested with either HpaII or HhaI (New England BioLabs), which are methylation sensitive restriction enzymes, in order to verify the methylation status of the methylated and mock methylated constructs prior to transfection.

### Cell culture, transient transfections, and dual luciferase assays

HEK 293 and GC-1spg cells were purchased from American Tissue Culture Collective (ATCC) (CRL-1573 and CRL-2053). The cells were grown in Dulbecco’s modified Eagles medium (DMEM) with high glucose (Cytiva) supplemented with 10% Fetal Bovine Serum (FBS) (Gemini Biosciences), 10 units/mL penicillin and streptomycin solution (Hyclone), 1x GlutaMAX glutamine supplement (Gibco), and 0.25 units/mL amphotericin B (Corning) in the presence of 5% CO_2_ at 37°C. For transfections, the cells were plated out at 50,000 cells/well for HEK 293 cells and 30,000 cells/well for GC-1spg cells in a 24-well tissue culture plate the day before transfection. For each transfection, 450 ng of the pCpGL firefly luciferase construct was transfected using PolyJet *in vitro* transfection reagent (SignaGen) according to manufacturer instructions. For all transfections, 50 ng of *Renilla* luciferase expression plasmid (pRL-TK) (Promega) was co-transfected for transfection efficiency normalization (500 ng of DNA total was added per transfection). Forty-eight hours post transfection, the cells were harvested and both the firefly and *Renilla* luciferase activities were measured from the same sample using the Dual-Luciferase Reporter Assay System (Promega) according to manufacturer instructions on a NOVOstar microplate reader (BMG Labtech). For each construct and treatment tested, four independent transfections were performed and assayed in duplicate. The firefly luciferase activity of each individual transfection was normalized against its *Renilla* luciferase activity by calculating the ratio of firefly to *Renilla* luciferase activity. Statistical significance was determined by calculating p values using a two-tailed unpaired student t-test.

## Results

Mammals have two SL9C genes (SLC9C1 and SLC9C2) whereas sea urchin possesses only one (http://bouzouki.bio.cs.cmu.edu/Echinobase/Search/SpSearch/) (Pedersen and Counillon, 2019; Windler et al., 2018). The sea urchin SLC9C protein shares nearly equal identity with both human SLC9C1 and SLC9C2 (NHE10 and NHE11) (Gardner and James, 2023). To ensure that each of the human, mouse, and rat genes we were comparing were truly all SLC9C1 orthologs, we first analyzed the chromosomal location and overall structure of these genes. For all three species, these genes reside in blocks of chromosomal synteny; the same upstream and downstream genes, in the same orientation, flank SLC9C1 in mouse, rat, and human (Figure 1A). Additionally, the SLC9C1 gene structure (exon-intron-exon) is very similar across mouse, rat, and human, mainly varying in intron lengths although the rat gene is predicted to contain one additional exon (Figure 1B).

**Figure 1.**
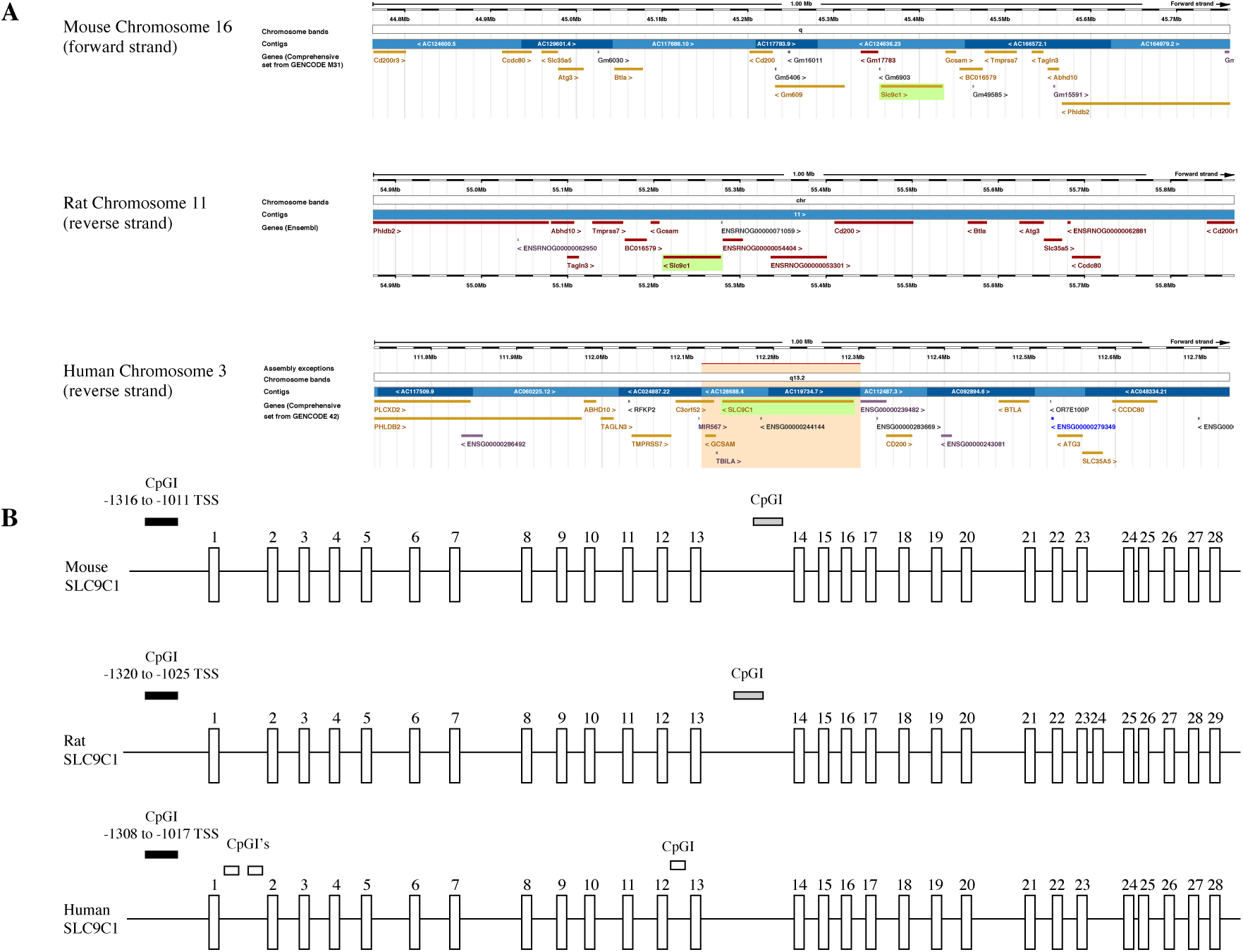
Evidence of Conservation of the SLC9C1 gene across mouse, rat, and human. A) The genes surrounding SLC9C1 (within 1.0 Mb) in mouse, rat, and human were analyzed using the Ensembl genome browser (Ensembl release 109) (Cunningham et al., 2022). There is clear evidence of synteny across the three species: the same flanking genes are present in all three species. The following flanking genes are found upstream of SLC9C1 in all three species: Cd200, Btla, Atg3, Slc35a5, and Cdc80. The following flanking genes are found downstream of SLC9C1 in all three species: Gcsam, Tmprss7, Tagln2, Abhd10, Phldb2. This suggests that SLC9C1 is evolutionarily conserved at the genomic level across these three mammalian species. The genomic coordinates for each SLC9C1 gene are mouse chromosome 16 (forward strand) 45,355,672 - 45,427,364, rat chromosome 11 (reverse strand): 55,211,792-55,278,285, and human chromosome 3 (reverse strand): 112,140,898-112,294,227. B) Schematic representations of the gene structures for mouse, rat, and human SLC9C1 are displayed (exons are represented by the open vertical boxes). Using DBCAT and SMS CpG island prediction software, a potentially conserved CpG island (CpGI) indicated by the darkened box was found in the same relative region about 1000 bp upstream of the transcription start site (TSS) in mouse, rat and human SLC9C1 (not to scale). The grey box indicates a less conserved CpGI identified in the intron between exons 13 and 14 in both mouse and rat SLC9C1 but not human SLC9C1. The unfilled boxes indicate CpGI’s identified in human SLC9C1 but not mouse or rat SLC9C1. The mouse gene is 71,735 bp long, the rat gene is 66,511 bp long, and human gene is 153,318 bp long.

Next, we identified potential promoters in the SLC9C1 genes using *in silico* software prediction. As expected, this software predicted promoters that covered the already annotated transcription start sites. The promoter sequences were aligned to each other in order to identify potentially conserved nucleotides (Figure 2A). The three species’ promoters exhibit high sequence conservation starting at ∼200 base pairs (bp) upstream of the transcription start site and continues past the transcription start site into the first exon. Interestingly, all three species possess a GC-box at the transcription start sites and also possess CCAAT boxes in the same location ∼50 bp upstream of the transcription start site (Figure 2A).

**Figure 2.**
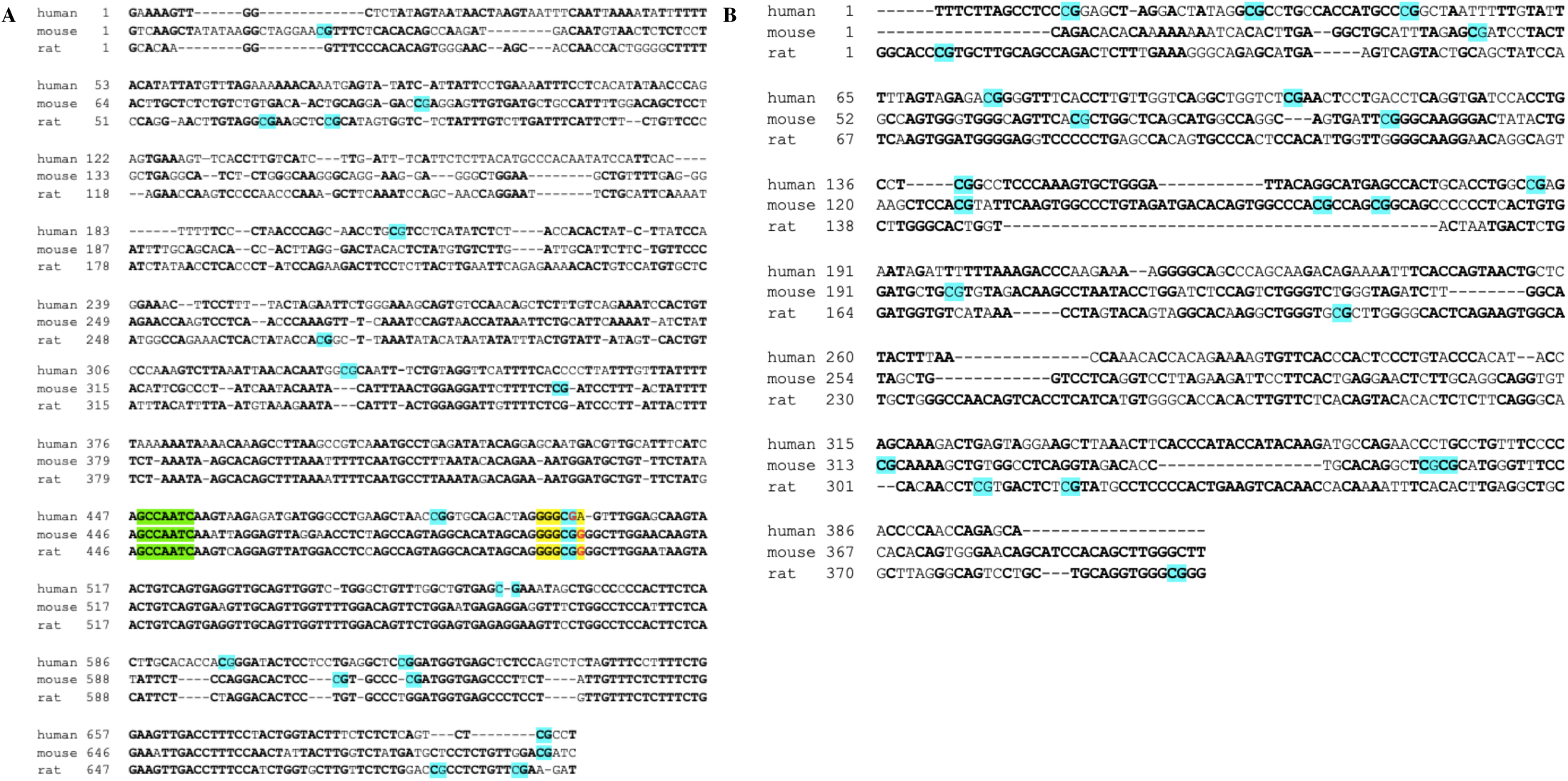
Sequence alignments of mouse, rat, and human SLC9C1 promoters and CpG islands. A) The predicted promoter regions (-500 to +200 relative to the TSS) are aligned across mouse, rat, and human SLC9C1 using T-Coffee. The nucleotides in bold lettering are identical to at least one other species. The red colored nucleotides indicate the transcription start sites (+1). The yellow highlighted nucleotides indicate a conserved GC box and SP-1 transcription factor binding site that is common to all three species in the same position at the transcription start site. The green highlighted nucleotides indicate a conserved CCAAT box and NF-Y transcription factor binding site that is in the same position in all three species ∼50 bp upstream of the transcription start site. Turquoise highlighted nucleotides are CpG dinucleotides. Note that there is no canonical TATA box observed in any of the three species’ promoters. The human and rat sequences share 56.65% identity, human and mouse 50.0% identity, and rat and mouse 67.26% identity. B) The conserved CpGI just upstream of the SLC9C1 promoter sequences (-1200 to - 800 of TSS) from mouse, rat, and human are aligned using T-Coffee. Turquoise highlighted nucleotides are CpG dinucleotides. The human and rat CpGI sequences share 35.59% identity, human and mouse 38.48% identity, and rat and mouse 48.83% identity. The nucleotides in bold lettering are identical to at least one other species.

We then identified potential CpG islands using software prediction. While there were several predicted CpG islands scattered throughout all three SLC9C1 genes, a particular CpG island stood out as being potentially conserved because it appeared at about 1000 bp upstream of the identified transcription start site of each of the three SLC9C1 genes (Figure 1B and 2B). Due to these CpG islands being so close to the transcription start site and located in the same relative position across the three species, these particular CpG islands as well as the predicted promoter regions across mouse, rat, and human SLC9C1 were chosen for analysis.

We next analyzed whether there is a difference in methylation of the SLC9C1 promoter and CpG island across the three mammalian species in testis versus somatic tissue (Figure 3). Bisulfite sequencing analysis of lung (a representative somatic tissue that does not express SLC9C1) and testis genomic DNA revealed a stark difference in methylation of the CpGI through the promoter region of the mouse SLC9C1 gene between the two tissues (87.5% methylation in lung and 2.8% in testis). There was a less obvious difference in methylation across this entire region (CpGI through the promoter) in both the rat (82.4% methylation in lung and 58.9% in testis) and in human (69.5% methylation in lung and 50.4% in testis) however, in both cases the methylation of this region was lower in the testis than it was in the lung (Figure 3).

**Figure 3.**
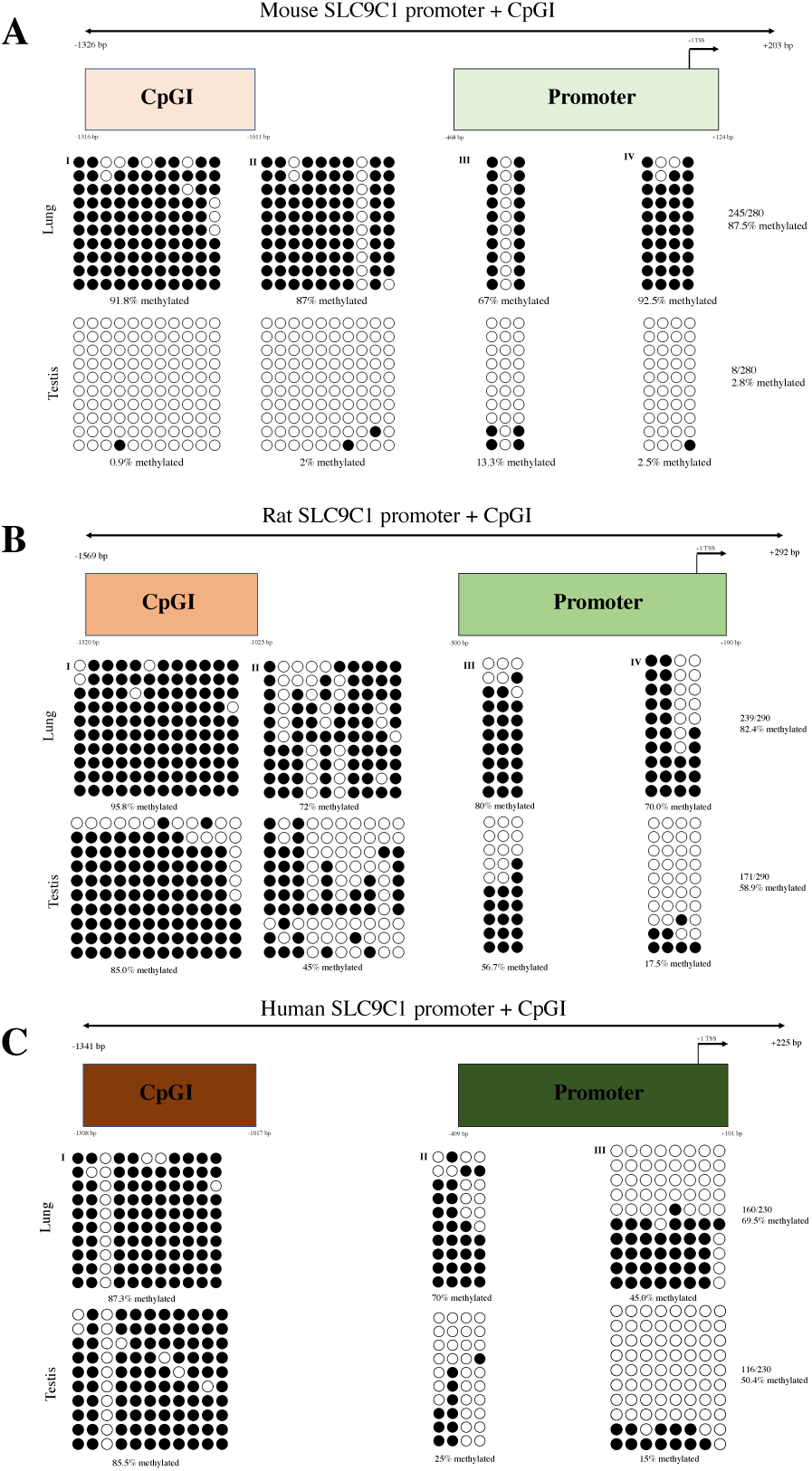
Bisulfite sequencing analysis of the mouse, rat, and human SLC9C1 promoters and CpG islands. A) Mouse lung and testis genomic DNA were bisulfite converted and then PCR amplified using 4 sets of primers displayed as four non-overlapping sub-regions I, II, III, IV with a total of 28 CpG dinucleotides. Sub-region I consists of nucleotides -1326 to -951 relative to TSS, sub- region II consists of nucleotides -915 to -511, sub-region III consists of nucleotides -431 to -45, and sub-region IV consists of -61 to +203. B) Rat lung and testis genomic DNA were bisulfite converted and then PCR amplified using 4 sets of primers displayed as four non-overlapping sub-regions I, II, III, IV with a total of 29 CpG dinucleotides. Sub-region I consists of nucleotides -1569 to -1097 relative to TSS, sub-region II consists of nucleotides -1025 to -554, sub-region III consists of nucleotides -624 to -230, and sub-region IV consists of -265 to +292. C) Human lung and testis genomic DNA were bisulfite converted and then PCR amplified using 3 sets of primers displayed as four non-overlapping sub-regions I, II, and III with a total of 23 CpG dinucleotides. Sub-region I consists of nucleotides -1341 to -960 relative to TSS, sub- region II consists of nucleotides -641 to -139, and sub-region III consists of nucleotides -260 to +225. Open circles represent unmethylated CpGs and closed circles represent methylated CpGs. Columns represent individual CpGs and rows represent individual clones. For each sample (lung and testis) a total of ten clones were analyzed by bisulfite sequencing.

By itself, the mouse SLC9C1 CpG island is clearly differentially methylated between lung and testis (sub-region I shows 91.8% methylation in lung and 0.9% in testis) (Figure 3). There is a small difference in methylation of the rat SLC9C1 CpG island (sub-region I is 95.8% methylated in rat lung and 85.0% in rat testis) but there is essentially no difference in methylation of the human SLC9C1 CpG island (87.3% methylated in human lung and 85.5% in human testis) between testis and lung (Figure 3).

The promoter regions of mouse, rat, and human SLC9C1 are differentially methylated (in mouse sub-region IV is 92.5% methylated in lung versus 2.5% in testis, in rat sub-region IV is 70.0% methylated in lung versus 17.5% in testis, and in human sub-region III is 45.0% methylated in lung versus 15.0% in testis) between the two tissues (Figure 3).

Our bisulfite sequencing data is consistent with publicly available methylome data (Li et al., 2018) (https://ngdc.cncb.ac.cn/methbank/search?item=&term=slc9c1) for mouse, rat and human SLC9C1 methylation across various tissues which suggests that both the mouse and rat SLC9C1 promoter is hypomethylated in sperm cells but hypermethylated in other tissue and the human SLC9C1 promoter is also less methylated in semen samples relative to the methylation profile of other somatic tissues.

We next tested the gene regulatory activity of the putative SLC9C1 promoters and CpG islands from the mouse, rat, and human genes using *in vitro* luciferase reporter assays. To do this, we cloned the regions-of-interest from the SLC9C1 genes upstream of the firefly luciferase open reading frame (ORF) into the pCpGL vector, a plasmid without any endogenous CpG dinucleotides. We cloned either the promoter (prom), the CpG island (CpGI), the whole region of the CpGI through the promoter (“inter” designating the intervening region between the CpG island and the promoter; CIP), or just the CpG island and promoter (without the intervening region; CP) into the pCpGL vector (Figure 4A). These constructs were then transfected into either HEK 293 cells as a representative somatic cell line or GC-1spg cells (Hofmann et al., 1992) as a representative male germ cell line and compared the luciferase activity to the same cells transfected with the promoter-less pCpGL construct (pCpGL-Basic). We found that all of the SLC9C1 promoters exhibited activity in both HEK 293 and GC-1spg cells although in all cases, the promoter activity was greater in HEK 293 than in GC-1spg cells (Figure 4B and 4C). The rat and human SLC9C1 promoters exhibited much greater activity than the mouse promoter in HEK 293 cells, while all three SLC9C1 promoters exhibited roughly equal activity in GC- 1spg cells (Figure 4B and 4C).

**Figure 4.**
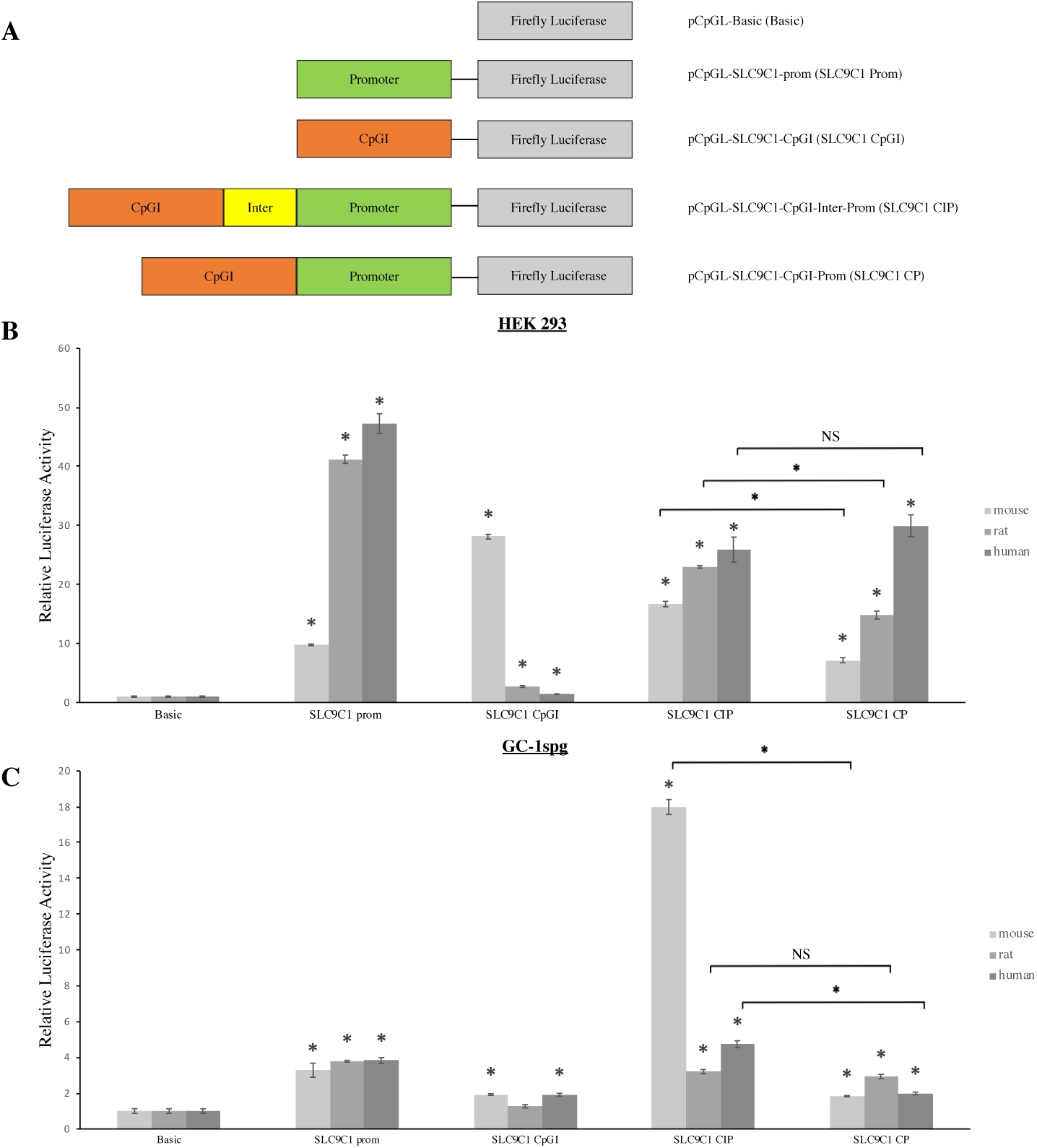
Promoter activity of mouse, rat, and human SLC9C1 putative promotors and CpG islands. A) Elements from the mouse, rat, or human SLC9C1 genes were cloned into the pCpGL base vector and the ability to drive the expression of firefly luciferase was assessed in both a somatic cell line (HEK293) and a spermatogonial cell line (GC-1 spg). For each species, reporter constructs containing just the promoter, just the upstream CpGI, the entire region containing the CpGI plus the promoter plus the region between them (inter) (CIP), and entire region minus the inter region (promoter plus the CpGI only) (CP) were tested for the ability to drive the expression of firefly luciferase. The promoter-less pCpGL-Basic construct was used to establish a baseline luciferase activity for the vector. B&C) Firefly luciferase activity in transfected HEK 293 cells (B) or GC-1spg cells (C). The firefly luciferase activity was normalized to *Renilla* luciferase activity from the co-transfected transfection efficiency control vector pRL-TK and is depicted as fold increase over the promoter-less pCpGL-Basic vector activity (set at 1.0). Note that all of the tested constructs displayed significant promoter activity in HEK 293 cells when compared to the pCpGL-Basic vector. For B) and C), (*) over bars indicates statistically significant difference with p values <0.05 from pCpGL-Basic and (*) over bracket indicates statistically significant difference with p values <0.05 between bars at each end of bracket. (NS) indicates not statistically significant.

Interestingly, the mouse CpG island by itself exhibited strong promoter activity in HEK 293 cells, while the human and rat CpG islands exhibited only modest promoter activity in this cell line; the mouse CpG island displayed 10 and 20 fold the promoter activity of the rat and human CpG islands respectively in HEK 293 cells (Figure 4B). In GC-1spg cells, the mouse and human CpG islands exhibited low, but detectible promoter activity whereas the rat CpG island displayed no activity in these cells (luciferase activity was not significantly different than in cells transfected with the promoter-less pCpGL-Basic; Figure 4C).

The entire CpG island through the promoter (CIP) regions from all three species exhibited strong promoter activity in HEK 293 cells (Figure 4B). The mouse CIP possessed the strongest promoter activity in GC-1spg cells, comparable to its activity in HEK 293 cells, while the promoter activity of the CIP regions from rat and human displayed low activity in these cells (Figure 4C). All of the constructs containing just the CpG island and promoter without the intervening sequence (CP) exhibited promoter activity in both HEK 293 and GC-1spg cells (Figure 4B and 4C). In HEK 293 cells, the promoter activity for the mouse and rat CP was lower than the promoter activity of the respective CIP for these same species whereas there was no difference between the promoter activities of the human CIP and CP (Figure 4B). In GC-1spg cells, the promoter activity of the mouse and human CIPs were greater than the activities of their respective CP regions while the rat CIP and CP regions displayed equivalent promoter activity in these same cells (Figure 4C).

Having established the promoter activity of each of these SLC9C1 DNA elements, we next tested whether the promoter activities of these elements were sensitive to DNA methylation. To do this, we *in vitro* methylated or mock methylated each construct and then transfected them into HEK 293 and GC-1spg cells (Figure 5). Any differences in promoter activity, because the pCpGL vector does not contain any endogenous CpG dinucleotides (Klug and Rehli, 2006), is due to methylation of the CpG dinucleotides in the SLC9C1 genomic fragments in the reporter constructs. As expected, the pCpGL-Basic vector did not display methylation-sensitive promoter activity in either cell line (Figure 5B and 5C). The SLC9C1 promoters each displayed a different pattern of sensitivity to DNA methylation: the mouse promoter was not sensitive to DNA methylation in either HEK 293 or GC-1spg cells, the rat promoter was methylation-sensitive (displayed reduced promoter activity when methylated compared to mock methylated) in both cell lines (although it was more sensitive in GC-1spg cells) and the human promoter was very methylation-sensitive in HEK 293 cells but not sensitive in GC-1spg cells (Figure 5B and 5C).

**Figure 5.**
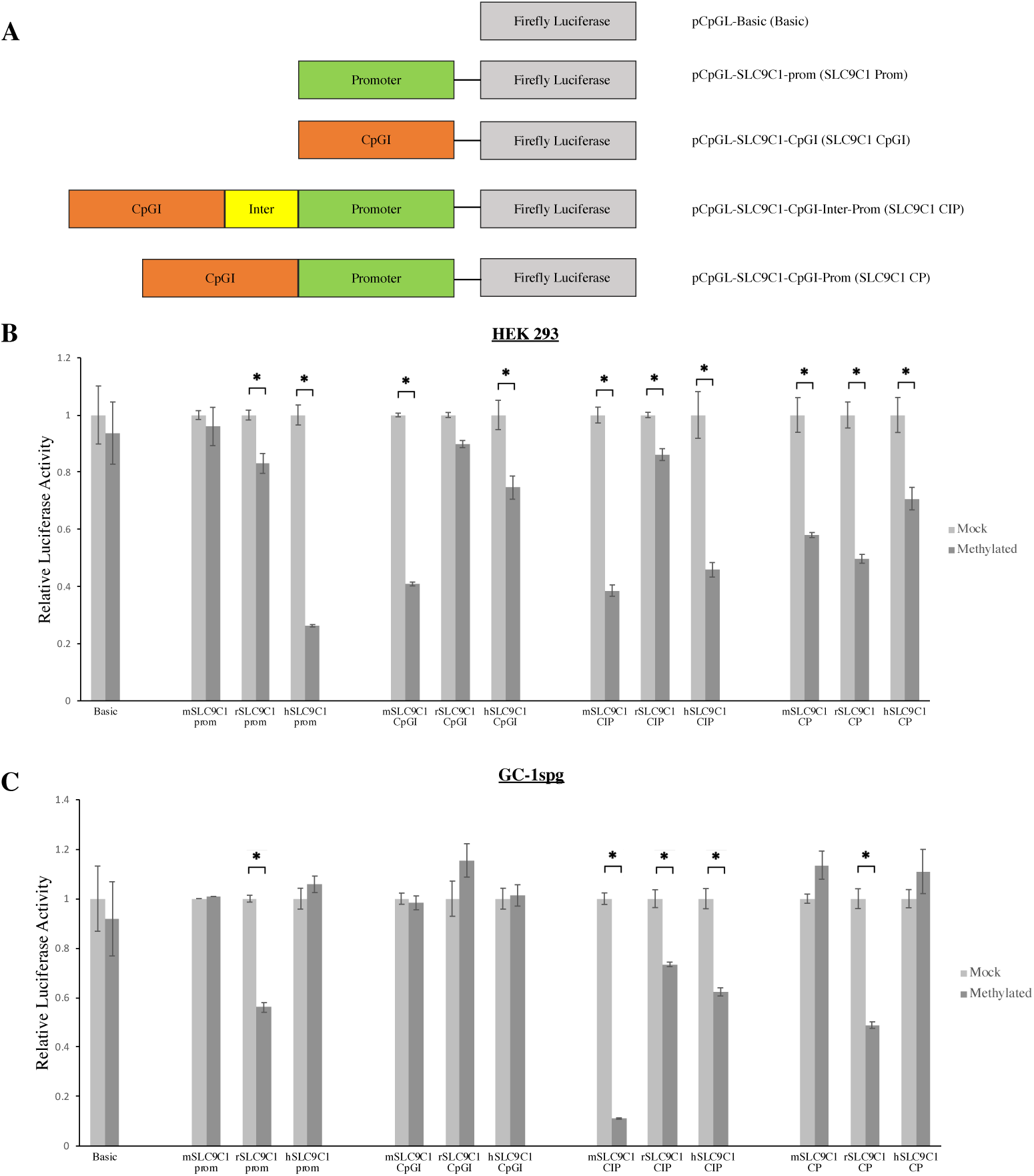
Methylation-sensitivity of mouse, rat, and human SLC9C1 promoters and CpG islands. A) Elements from the mouse, rat, or human SLC9C1 genes were cloned into the pCpGL base vector and the ability to drive the expression of firefly luciferase was assessed in both a somatic cell line (HEK293) and a spermatogonial cell line (GC-1 spg). For each species, reporter constructs containing just the promoter, just the upstream CpGI, the entire region containing the CpGI plus the promoter plus the region between them (inter) (CIP), and entire region minus the inter region (promoter plus the CpGI only) (CP) were tested. The reporter constructs were *in vitro* M. SssI methylated or mock methylated and then transfected. B&C) The sensitivity of the promoter activity to CpG methylation in HEK 293 cells (B) or GC-1spg cells (C). The firefly luciferase reporter activity (normalized to *Renilla* luciferase activity) of the methylated constructs are shown as the fold change compared to the mock methylated constructs (set at 1.0). Error bars indicate mean ± SEM of four independent transfections performed in duplicate. For B) and C), (*) over brackets indicates statistically significant difference with p values <0.05 between bars at each end of the bracket.

The CpG islands upstream of the SLC9C1 gene from mouse and human both displayed methylation-sensitive promoter activity in HEK 293 cells while the rat CpG island did not (Figure 5B). In contrast, the promoter activity of the CpG islands in all three species were insensitive to methylation in GC-1spg cells (Figure 5C).

For all three species, the entire CIP region displayed methylation-sensitive activity in both cell types (Figure 5B and 5C) with the mouse CIP displaying a reduction of 89% of promoter activity in GC-1spg cells when methylated (Figure 5C). Removing the sequence between the CpG islands and their respective promoters (the CP constructs) resulted in little change in HEK 293 cells: both the CP and CIP constructs displayed methylation-sensitive promoter activity (Figure 5B) however removal of the intervening region appears to increase methylation-sensitivity for the rat construct (Figure 5B). In GC-1spg cells, loss of the intervening region eliminates the methylation-sensitivity of the promoter activity for both the mouse and human constructs but has no effect on methylation-sensitive promoter activity of the rat construct (Figure 5C).

One of the most compelling findings from our analyses to this point was that the mouse SLC9C1 CpG island and promoter were almost completely methylated in lung and almost completely demethylated in testis. This, combined with the mouse CpG island’s strong promoter activity by itself in HEK 293 cells (that was sensitive to methylation) as well as the robust and methylation-sensitive promoter activity of the mouse CIP in both HEK 293 and GC-1spg cells, prompted us to examine the effect that the mouse CpG island had on the methylation-sensitive and -insensitive promoter activity of the rat and human inter-promoter (IP) regions. Toward that end, we assembled reporter constructs in which the rat and human CpG island were replaced with the mouse CpG island, leaving intact the rest of the rat or human IP fragments (Figure 6A). We transfected these chimeric constructs into HEK 293 and GC-1spg cells to determine the influence that the mouse CpG island has on these other species’ regulatory elements (Figure 6B- E).

**Figure 6.**
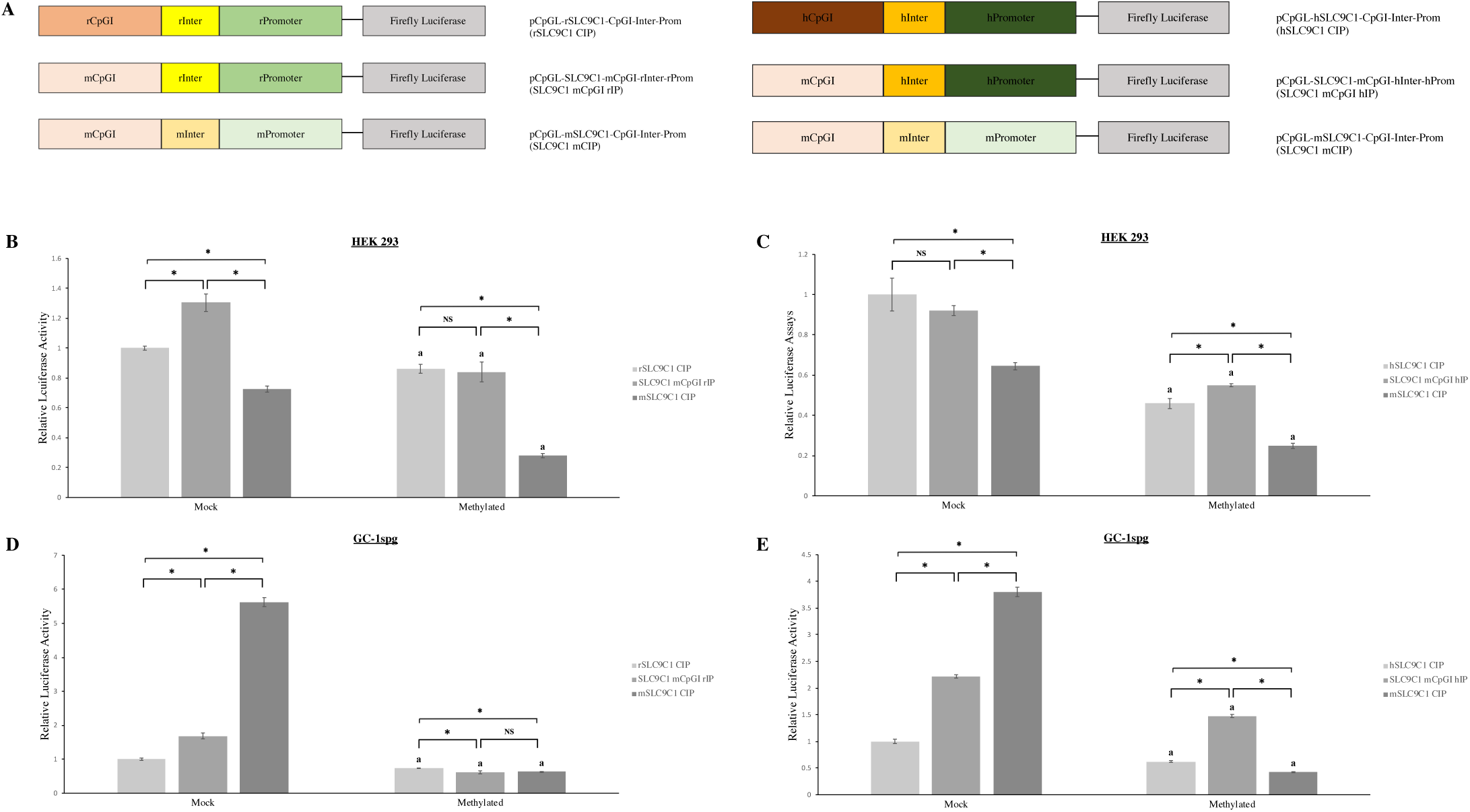
Influence of the mouse SLC9C1 CpG island on the activity of the rat and human regulatory elements. A) Schematic representations of the chimeric SLC9C1 luciferase reporter constructs. Reporter constructs in which the mouse CpG island replaced either the rat or human CpG island in the respective CpGI-inter-promoter (CIP) regulatory regions were assembled and transfected into HEK 293 and GC-1 spg cells. B&C) The promoter activity of the chimeric constructs consisting of the mouse SLC9C1 CpGI cloned upstream of either the rat (B) or human (C) inter-promoter region was analyzed in HEK 293 cells. D&E) The promoter activity of the chimeric constructs consisting of the mouse SLC9C1 CpGI cloned upstream of either the rat (D) or human (E) inter- promoter regions were analyzed in GC-1 spg cells. The reporter activity of mock methylated or methylated chimeric constructs are normalized to the activity of the mock methylated or methylated native human (B&D) or rat (C&E) SLC9C1 CIP promoter activity (set to 1.0). Error bars indicate mean ± SEM of four independent transfections performed in duplicate. (a) indicates this methylated construct exhibits a statistically significant difference with p values <0.05 from its respective mock methylated construct. (*) over brackets indicates statistically significant difference with p values <0.05 between bars at each end of the bracket.

Swapping out the rat CpG island for the mouse CpG island in the rat CIP increased promoter activity for the mock methylated constructs in both cell lines (Figure 6B and 6D). The promoter activity of the mouse-rat chimeric construct was significantly more sensitive to methylation in the GC-1 cells than the native rat CIP construct (Figure 6D) but the sensitivity to methylation in HEK 293 cells was equivalent to the native rat CIP construct (Figure 6B). The mock methylated mouse-human chimeric construct displayed increased promoter activity compared to the native human CIP in GC-1spg cells but not in HEK293 cells (Figure 6C and 6E). The mouse-human chimeric construct appears to be less sensitive to methylation than the native human CIP in HEK 293 cells (Figure 6C) but has an equivalent sensitivity to methylation in GC-1spg cells (Figure 6E). When methylated, the mouse-human chimeric CIP possess increased promoter activity compared to the methylated native human CIP in both cell lines (Figure 6C and 6E).

Next, because the mouse CpG island could influence the promoter activity of the rat and human CIPs, we tested its influence on other promoters by generating luciferase reporter constructs that placed the mouse CpG island upstream of either the minimal CMV promoter or the human EF1α promoter (and first intron) (Figures 7 and 8) and assessed the ability of these constructs to drive luciferase expression, in both the methylated and unmethylated states, in HEK 293 and GC-1spg cells. The constructs were mock methylated or *in vitro* methylated, transfected into either HEK or GC-1spg cells, and luciferase activities, relative to the CMV promoter or EF1α promoter alone, were determined.

**Figure 7.**
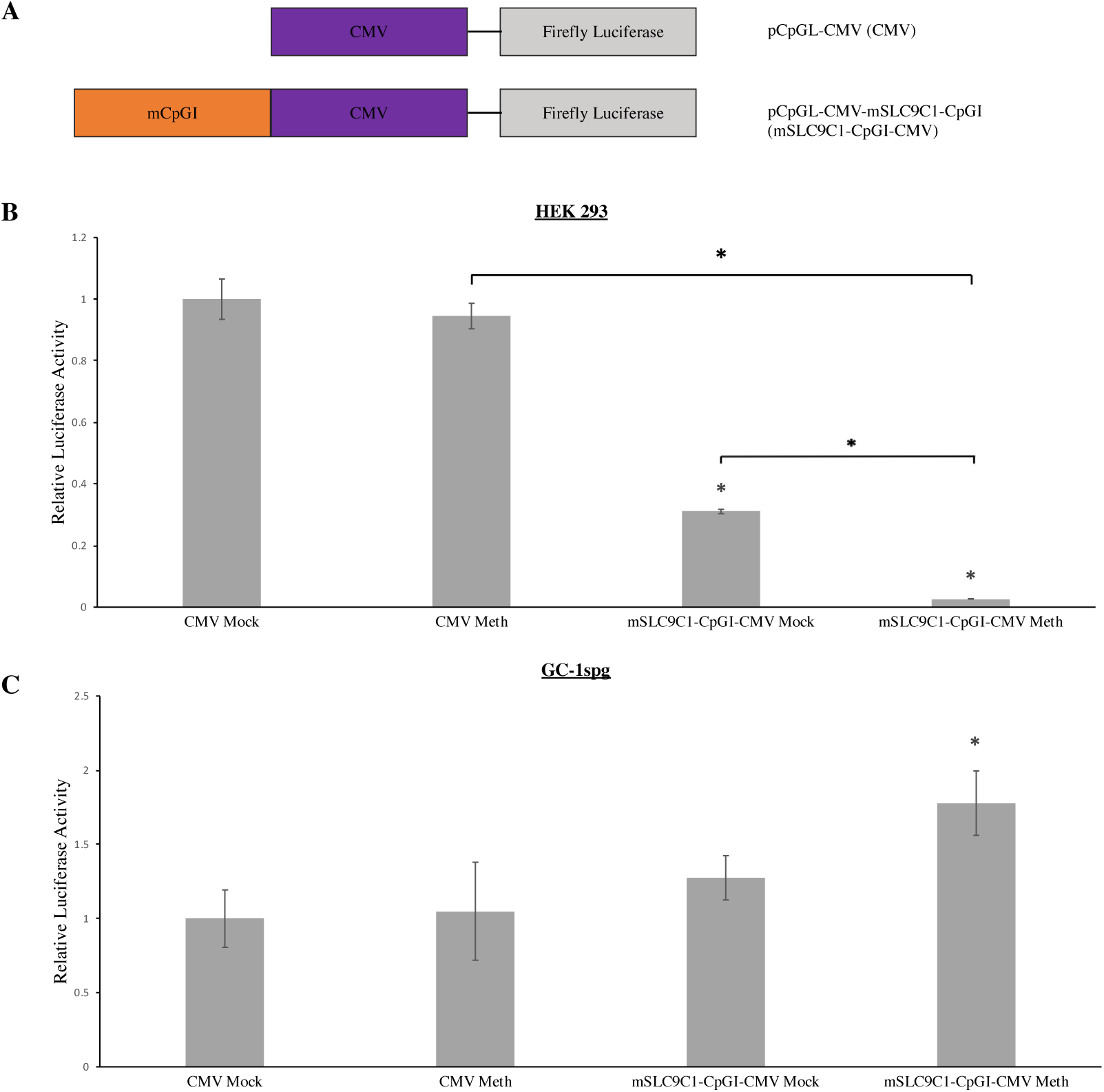
Influence of the mouse SLC9C1 CpG island on the activity of the CMV promoter. A) A schematic depicting luciferase constructs containing the mouse SLC9C1 CpGI cloned upstream of the minimal CMV promoter. B) The reporter activity of mock methylated or methylated chimeric constructs were normalized to the activity of the mock methylated CMV- only promoter activity (set to 1.0) in HEK 293 cells. C) The reporter activity of mock methylated or methylated chimeric constructs were normalized to the activity of the mock methylated CMV- only promoter activity (set to 1.0) in GC-1 spg cells. Error bars indicate mean ± SEM of four independent transfections performed in duplicate. (*) over brackets indicates statistically significant difference with p values <0.05 between bars at each end of the bracket.

**Figure 8.**
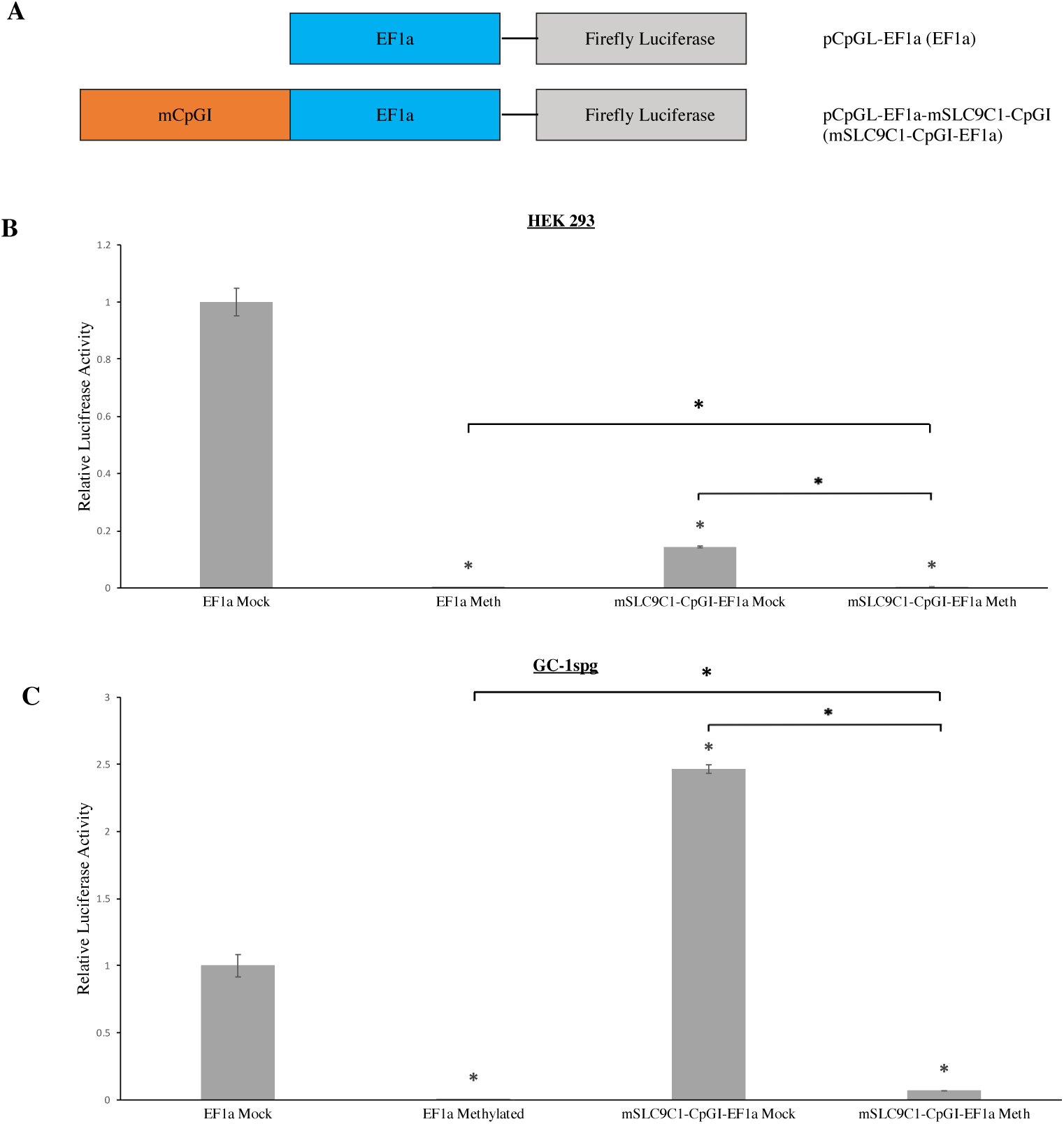
Influence of the mouse SLC9C1 CpG island on the activity of the EF1α promoter. A) A schematic depicting luciferase constructs containing the mouse SLC9C1 CpGI cloned upstream of the EF1α promoter. B) The reporter activity of mock methylated or methylated chimeric constructs were normalized to the activity of the mock methylated EF1α-only promoter activity (set to 1.0) in HEK 293 cells. C) The reporter activity of mock methylated or methylated chimeric constructs were normalized to the activity of the mock methylated EF1α-only promoter activity (set to 1.0) in GC-1 spg cells. Error bars indicate mean ± SEM of four independent transfections performed in duplicate. (*) over brackets indicates statistically significant difference with p values <0.05 between bars at each end of the bracket.

The CMV promoter does not contain any CpG dinucleotides and is therefore insensitive to methylation in either cell line (Figure 7B and 7C). In HEK 293 cells, the mouse CpG island reduced the activity of the CMV promoter in the unmethylated state (Figure 7B) but had no effect on CMV promoter activity in GC-1spg cells (Figure 7C). Therefore, the mouse CpG island has a silencer effect on the CMV promoter in the somatic cell line but does not influence the activity of the same promoter in the spermatogonial cell line. When methylated, the promoter activity of the mouse CpG island-CMV promoter was lost in HEK 293 cells but was significantly increased in GC-1spg cells (Figure 7B and 7C). Thus, when the mouse SLC9C1 CpG island was *in vitro* methylated it had a very strong silencer effect on the CMV promoter in HEK 293 cells but had an enhancer effect in GC-1spg cells (Figure 6B and 6C).

Containing multiple CpG dinucleotides, the EF1α promoter by itself is exquisitely sensitive to methylation in both HEK 293 and GC-1spg cells; when methylated, the activity of the EF1α promoter is essentially completely lost in both cell lines (Figure 8B and 8C). The mouse CpG island-EF1α promoter displayed reduced activity compared to the EF1α promoter alone in HEK 293 cells (Figure 8B) and increased activity compared to the EF1α promoter alone in GC-1spg cells (Figure 8C). In HEK 293 cells, the mouse CpG island-EF1α promoter essentially lost all activity when methylated (Figure 8B and 8C). In GC-1spg cells, the mouse CpG island-EF1α promoter lost most, but not all, promoter activity when methylated (Figure 8B and 8C). The residual activity of the methylated CpG island-EF1α could represent promoter activity driven by the CpG island alone, as it is insensitive to methylation in these cells (Figure 5C).

## Discussion

Numerous previous studies have demonstrated the importance of proper expression of SLC9C1 (NHE10) in testis/sperm for maintaining male fertility across many mammalian species. A study in bull sperm found that SLC9C1 mRNAs were upregulated in sperm considered to be good osmoregulators as compared to poor osmoregulators (Lavanya et al., 2021), highlighting the potential importance of increased SLC9C1 gene expression in maintaining fertilizing capacity in mammals. Additionally, SLC9C1 was found to be a downregulated differentially expressed gene in non-obstructive azoospermia (NOA) patients (Hu et al., 2021). SLC9C1 has been shown to be essential for male fertility in both mice (Wang et al., 2003) and humans (Cavarocchi et al., 2021). Additionally, reduced levels of NHE10 protein expression was found in a group of subfertile human male patients (Zhang et al., 2017). This particular study is of interest because it reveals a correlation between NHE10 protein levels and male fertility, however it does not investigate what the mechanism(s) of the reduced NHE10 protein expression in these subfertile patients was. Further work is needed to see if aberrant DNA methylation, such as hypermethylation of the SLC9C1 promoter, is one of the causes of the decreased NHE10 expression in these human male patients. All of the above work is strong evidence that the proper expression of SLC9C1 (NHE10) is essential for male fertilizing ability across mammalian species.

The overall goals of these studies were to begin to identify the DNA methylation-dependent and -independent regulatory mechanisms involved in the tissue-specific expression of the SLC9C1 gene in mice, rats, and humans. Here we report the identification and characterization of two DNA regulatory elements, the promoter and a CpG island located ∼1000bp upstream of the TSS, that are shared by all three mammalian SLC9C1 genes. Although both of these 5’ cis elements are present in the genes for all three species, our analyses suggest that they display some species-specific differences in gene regulatory activity in both the methylated and unmethylated states.

Once we confirmed, using sequence comparisons, syntenic analysis, and predicted gene structure that the genes from all species were SLC9C1 homologs (Figure 1), our *in silico* analysis identified putative promoters as well as a common upstream CpG island in the genes for each species (Figures 1 and 2). Further sequence analysis of the putative promoters demonstrated a high degree of sequence identity (higher identity between rat and mouse than between either rodent and human) and uncovered two conserved DNA regulatory sequences shared by all three SLC9C1 gene promoters: a GC box overlapping with the transcription start site and a CCAAT box ∼50 base pairs upstream of the transcription start site (Figure 2A).

The GC box is a highly conserved DNA sequence motif “GGGCGG” which the Sp1 transcription factor has been well characterized to bind to where it functions to recruit RNA polymerase II in order to initiate transcription (Kadonaga et al., 1986). One study using the human metallothionein (hMT) IIA gene promoter as a model found that CpG methylation of the CpG dinucleotide in the endogenous GC box of this gene promoter did not inhibit SP1 transcription factor binding in *in vitro* DNA-protein binding assays (Harrington et al., 1988).

However, another study using the human *α*_1d_ -andrenergic receptor gene promoter as a model found that methylation of promoter constructs blocked SP1 activation of gene expression using luciferase reporter assays and also found that, in cell lines expressing various levels of *α*_1d_- andrenergic receptor RNA, expression levels were inversely correlated with methylation of the GC boxes in the promoter (Michelotti et al., 2007). These later results suggest that GC box methylation may play a role in regulating SP1 transcription factor binding in a cell/tissue-specific context and suggest that further analysis into the role that methylation plays in SP1 regulation of SLC9C1 expression is needed.

The CCAAT-box is a common regulatory element of eukaryotic promoters that binds to the NF-Y protein (also known as CBF) and is normally found around -60 to -100 bp upstream of the transcription start site (Mantovani, 1999). In all three studied SLC9C1 genes, the CCAAT-box is ∼50 bp upstream of the transcription start site (Figure 2A). The CCAAT box is often critical for initiating RNA polymerase recruitment in genes that lack a canonical TATA box (Mantovani, 1999), which all three of these mammalian SLC9C1 genes lack. Binding of NF-Y to the CCAAT-box is thought to configure the DNA in such a way that other transcription factors recognizing that sequence are able to bind (Mantovani, 1999), thus NF-Y acts as a master regulator for more gene-specific transcription factor binding. Future experiments into the specifics of the CCAAT-box and NF-Y binding in the control of mammalian SLC9C1 gene regulation are also called for.

Having identified the putative promoters and the common upstream CpG island, we used bisulfite sequencing to determine whether these were differentially methylated regions in the genes from these three species. It should be noted that while SLC9C1 appears to be most highly expressed in spermatocytes and spermatids in mouse and human testis (https://orit.research.bcm.edu/MRGDv2), these are only a subset of the cells found in the testis.

Further, even genes expressed in only specific cell types of the testis have been shown to be differentially methylated across these cells in the same testis tissue (Ben Maamar et al., 2022). In our bisulfite sequencing analyses, we used the whole testis and therefore our results represent the average of the SLC9C1 methylation pattern in all the cells of the testis (and the lung).

In the mouse, the SLC9C1 promoter, the CpG island, and the intervening regions are almost completely methylated in the lung but nearly totally unmethylated in the testis (Figure 3C), suggesting that DNA methylation of these regulatory elements may be involved in repressing expression in somatic tissues. On the other hand, while the uniform demethylation of these regions in all/most cell types in the testis may prime this region for gene activation, the expression of cell type-specific transcription factors and/or other epigenetic modifications would be needed to drive the cell-specific expression of the mouse SLC9C1 gene during spermiogenesis. In rat and human there is a less pronounced difference in overall methylation of the SLC9C1 promoter, the CpG island, and the intervening region when comparing the lung to the testis, however a measurable hypomethylation is still noted in the testis (Figure 3B and 3C). Although our luciferase reporter assays revealed that some of these gene regulatory sequences are methylation sensitive (discussed below), this suggests, unlike in the mouse where we see near complete demethylation in the testis, that these SLC9C1 regions may be only hypomethylated in the subset of specific cells in the rat and human testis that actually express the gene.

Interestingly, while the total number of CpG dinucleotides in the CpG island, the intervening sequence, and the promoter are similar between the three species (28 CpGs for mouse, 29 CpGs for rat, and 23 for CpGs human), the CpG dinucleotides are differentially distributed in the rodent genes compared to the human: the human gene lacks CpG dinucleotides in the intervening region and has a higher density of CpG dinucleotides closer to the transcriptional start site than do the rodent genes (Figure 3). The differential location of the CpG dinucleotide densities suggests a potential shift of methylation-sensitive DNA regulation from the CpG island towards the transcription start site for the human gene.

As noted, there is clear evidence of primary sequence conservation of the SLC9C1 promoters from these three species (Figure 2A) and we see similar differential methylation patterns of the promoters (Figure 3). On the other hand, there is a difference in the methylation pattern in rat and human CpG islands compared to the mouse (Figure 3) and these CpG islands share little sequence conservation (Figure 2B). These data are in agreement with the findings that whole genome tissue-specific methylation across the mouse, rat, and human genomes were similar in all three species mainly in areas of primary DNA sequence conservation like we find in these promoters, probably due to the conservation of common transcription factor binding sites (Zhou et al., 2017).

Having identified prospective cis-regulatory elements, some of which displaying the potential for DNA methylation dependence based on the presence of differential methylation in somatic tissue vs the testis, we next characterized their gene regulatory activity using reporter assays in cultured cells. Our results suggest that the SLC9C1 cis-regulatory elements from these species drive different methylation-sensitive and methylation-insensitive gene regulatory activity in somatic vs. male germ cells.

The activity of the human promoter was very sensitive to methylation in HEK 293 cells but insensitive in GC-1spg cells (Figure 5B and 5C) suggesting that DNA methylation of this region would inhibit SLC9C1 expression in somatic cells but not in male germ cells. Like the promoter, DNA methylation inhibited promoter activity of the human CpG island in HEK 293 cells but not in GC-1spg cells (Figure 5B and 5C). Whether separated by the intervening sequence or not, the human promoter and CpG island were methylation sensitive in HEK 293 cells suggesting that methylation of these regions in somatic tissues could be a mechanism to suppress unwanted SLC9C1 expression outside of the testis. Although the human promoter and CpG island promoter activity were individually insensitive to methylation in the male germ cells, together in context with the intervening sequence display methylation sensitivity that was lost when the intervening sequence was removed (Figure 5B and 5C). It is unclear how the intervening sequence confers this methylation sensitivity as there are no CpG dinucleotides in this region of the human SLC9C1 gene, perhaps the orientation of the human promoter and CpG island separated by the intervening sequence positions the CpG dinucleotides in these two regions such that looping of the DNA creates interactions that confer methylation sensitivity that are not present individually or when they are immediately adjacent to each other. In any case, DNA methylation of the human promoter and CpG island would make an effective mechanism for suppressing SLC9C1 gene expression in somatic tissue and, if oriented properly, in the SLC9C1 non-expressing cells of the testis. Overall, our data suggests that in HEK 293 cells, the human promoter is what is driving most of the methylation-sensitive gene regulatory activity, which correlates with our bisulfite sequencing data where in the lung, the human promoter region is clearly more methylated than it is in the testis (45.0% vs 15.0% methylated) (Figures 3C and 5B).

Like the human promoter, the rat SLC9C1 promoter displays strong activity in HEK 293 cells and displays equivalent activity to the other species’ promoters in GC-1spg cells (Figure 4b and 4C). The rat CpG island displays modest activity in the somatic cells but no detectible activity in the GC-1spg cells (Figure 4B and 4C). The CpG island and the promoter in context with the intervening sequence (CIP) are active in HEK-293 cells and that activity is reduced when the intervening sequence is removed (CP; Figure 4B). In GC-1spg cells, the rat CIP displays modest activity but unlike the human, that activity is unchanged when the intervening sequence is removed (Figure 4C). In addition, only the rat reporter constructs that contain the promoter show sensitivity to methylation and that sensitivity is not influenced by the presence or absence of the intervening sequence in either cell type (Figure 5B and 5C) even though multiple CpG dinucleotides are present in the rat intervening sequence (Figure 3B). Taken together, these results point to the rat promoter (like the human promoter) as the most important element of those tested for driving SLC9C1 expression in both somatic and male germ cells. In addition, it appears that methylation of the promoter is a potential mechanism to repress the expression of the rat SLC9C1 gene in somatic cells and in cells of the testis in which its expression is not required. This functional data is consistent with our findings that the largest difference in differential methylation between rat testis and lung is in the promoter, particularly in the CpG dinucleotides closest to the transcriptional start site (Figure 3B).

Compared to the human and rat promoters, the mouse promoter displayed much lower activity in HEK-293 but in GC-1spg cells the mouse promoter activity was nearly identical to the human and rat activity (Figure 4B and 4C). On the other hand, the mouse CpG island displayed very strong promoter activity by itself in HEK-293 cells, much stronger than the minimal promoter activity of the rat and human CpG islands and much stronger than the activity of the mouse promoter itself (Figure 4B). Although weak compared to its activity in HEK-293 cells, the mouse CpG island did display promoter activity in GC-1spg cells (Figure 4B). When the mouse promoter and CpG island are in the same construct, they display high (and equivalent) promoter activity in both cell lines when separated by the intervening sequence (the CIP construct; Figure 4B and 4C) but lose much of that activity when the intervening sequence is eliminated (the CP construct; Figure 4B and 4C). This suggests that sequences within the intervening sequence or the spacing of the promoter and the CpG island with respect to each other is important for promoter activity.

The activity of the mouse SLC9C1 promoter was not found to be methylation-sensitive in either cell line tested whereas the promoter activity of the mouse CpG island was strongly inhibited by methylation in the somatic cell line but not in the male germ cell line (Figure 5B and 5C). The mouse CIP promoter activity is methylation sensitive in both cell lines but removing the intervening sequence eliminates the methylation sensitivity in the male germ cell line only (Figure 5B and 5C). Taken together, these data suggest that, of the elements tested, it is likely the mouse SLC9C1 CpG island that is playing the most important role in preventing SLC9C1 expression in somatic cells: when methylated, as in the case of somatic tissue such as lung, aberrant expression is suppressed. On the other hand, both the mouse promoter and the CpG island individually are insensitive to methylation in GC-1spg cells however, when present together with the intervening sequence, the expression becomes very sensitive to methylation (Figure 5C). These results suggest that methylation of CpG dinucleotides in the intervening region between the CpG island and the promoter is a mechanism by which SLC9C1 expression could be shut down in cells of the testis that do not express this gene and that methylation in this region is more important than methylation in the promoter or CpG island for this regulation. The overall lack of methylation of the CpG dinucleotides in this region (as well as the CpG island and the promoter) that we see in the mouse testis (Figure 3A), would provide a permissive state for SLC9C1 expression in the appropriate testicular cells.

Having found that the mouse CpG island had a strong influence on the activity of the regulatory elements present in the 5’ region of its own gene, we wanted to examine if the mouse CpG island was able to modify the activity of the regulatory elements of the other two species’ genes. Towards that end, we generated chimeric constructs in which the native CpG island of the human and rat constructs were replaced by the mouse CpG island (Figure 6A).

Whether methylated or unmethylated, the chimeric construct in which the human CpG island was replaced with the mouse CpG island displayed promoter activity more like the human CIP than the mouse CIP in HEK 293 cells (Figure 6C) demonstrating that the mouse CpG island does not influence the activity of the human promoter in this somatic cell line. These results support our findings that the human promoter is the most important element (of those we examined) for the methylation-sensitive and -insensitive promoter activity of the human gene regulatory elements in HEK 293 cells and that this activity is influenced very little by its own CpG island or by the CpG island from the mouse gene. In GC-1spg cells, when the human CpG island is replaced by the mouse CpG island, the promoter activity increases to a level intermediate between that of the native human (lower) and mouse (higher) construct activities (Figure 6E). Further, promoter activity of the chimeric mouse/human construct displayed methylation sensitivity in line with the native human construct: the native human and mouse/human chimeric constructs were less sensitive to loss of activity due to methylation than the native mouse construct in GC-1spg cells (Figure 6E). Taken together, these results confirm that the mouse CpG island enhances promoter activity when in context with an intervening sequence and a promoter and does not influence methylation sensitivity in GC-1spg cells – the acute methylation sensitivity of the native mouse CIP is dependent on the presence of the mouse intervening sequence.

The mouse/rat chimeric CIP construct displayed promoter activity that was greater than both the rat and mouse native constructs in HEK 293 cells (Figure 6B) and, like the mouse/human chimeric construct, this activity displayed methylation sensitivity equivalent to the native rat construct (Figure 6D). Thus, in the somatic cell line, the mouse CpG island produced very similar outcomes in the rat construct as it did in the human construct: the mouse CpG island enhanced the promoter activity when unmethylated but did not influence promoter activity when methylated. Also like in the mouse/rat chimeric construct, replacing the rat CpG island with the mouse CpG island increased the promoter activity compared to the native rat CIP construct in the GC-1spg cells and this activity was lower than the native mouse CIP construct. Unlike the mouse/human chimeric construct which displayed higher residual promoter activity when methylated in GC-1spg cells, the methylated mouse/rat chimeric construct displayed low promoter activity, equivalent to the native CIP constructs (Figure 6D). Consistent with our findings using the mouse/human constructs, the mouse CpG island enhances the rat promoter activity in both cell types when unmethylated but unlike our findings with the mouse/human constructs, it does not alter promoter activity in GC-1spg cells when methylated. These results confirm that, while the mouse CpG islands can act as an enhancer, the most important cis-regulatory element, of those we tested, for regulating the expression of the rat and the human SLC9C1 genes are their respective promoters.

Having identified the potential modifier activity of the mouse CpG island using rat and human SLC9C1 cis-regulatory elements, we next examined whether this gene regulatory activity could influence gene expression from two unrelated promoters. Our luciferase reporter experiments demonstrate that the ability of the mouse CpG island to modify promoter activity depends on whether the constructs are expressed in a somatic or germ cell context. In HEK 293 cells, the unmethylated mouse CpG island significantly reduces expression from both the CMV and EF1α promoters (Figures 7B and 8B). When methylated, the promoter activity from the constructs containing the mouse CpG island is further reduced in these somatic cells (minimal activity for the CMV promoter and undetectable activity for the EF1α promoter; Figures 7B and 8B). On the other hand, the unmethylated mouse CpG island not only doesn’t inhibit expression in the GC-1spg cells from either promoter, it actually enhances expression from the EF1α promoter (Figures 7C and 8C). The mouse CpG island increased expression when either mock methylated or methylated in GC-1spg cells when placed upstream of either the CMV or EF1α promoter (Figures 7 and 8) which suggests that the mouse CpGI island upstream of either of these promoters enhances expression due to the transcription factors and relative epigenetic environment of these spermatogonial cells. This again points to the mouse SLC9C1 upstream CpG island as a DNA regulatory element that drives/enhances gene expression in germ cells but not in somatic cells, even when combined with irrelevant promoters. In contrast, the mouse SLC9C1 CpG island appears to exhibit a silencing effect on both the CMV and EF1α promoters in HEK 293 cells when mock methylated. Again, this is what would be expected if the CpGI acts to limit SLC9C1 expression to only germ cells. It appears that the SLC9C1 CpG island can exhibit either a silencing or enhancing effect depending on its methylation state and whether it is in a somatic or male germ cell environment.

It should be noted that our study only looked at the role of one specific epigenetic mechanism, DNA methylation, on regulating SLC9C1 gene expression. However future studies may examine how other epigenetic mechanisms such as histone modifications may be regulating tissue-specific SLC9C1 expression. It will be interesting to determine, in future studies, whether any of these other epigenetic mechanisms are conserved across these mammalian species in a similar way that methylation appears to be regulating SLC9C1 *in situ*.

Our work presented here provides the framework for future studies that utilize targeted epigenetic modifications of the SLC9C1 gene to modulate its expression either in cell culture or *in vivo*. With the advent of the CRISPR/Cas9 based systems to perform epigenetic modifications, it is now possible to perform targeted DNA methylation/demethylation *in vitro*. Our data suggests that if one wanted to increase or decrease SLC9C1 gene expression in a mouse cell, targeted demethylation or methylation of either the SLC9C1 CpG island and/or the intervening region would modulate gene expression. However, if one wanted to perform these targeted DNA methylation modifications to modulate SLC9C1 gene expression in a rat or human cell, one should probably only target the SLC9C1 promoter and not the CpG island, because the CpG island does not seem to have as much of a methylation-sensitive effect on gene expression in these species.

Future targeted methylation/demethylation studies of the SLC9C1 gene will reveal which specific CpG dinucleotides are responsible for regulating the observed DNA methylation-sensitive SLC9C1 gene expression. With the constant advances in gene therapy, it is possible that a currently infertile male with aberrant DNA hypermethylation of the SLC9C1 promoter, resulting in decreased NHE10 protein expression, could have their fertility restored by performing targeted DNA methylation of their SLC9C1 promoter to restore proper NHE10 protein expression levels. Additionally, it may be possible to decrease NHE10 protein expression in testis/sperm by performing targeted DNA methylation of the SLC9C1 promoter as a male contraceptive approach. DNA methylation is a reversible epigenetic modification and therefore targeted demethylation of the artificially induced methylation would restore normal protein expression, potentially acting as a reversible male contraceptive.

## Conclusion

Our study provides interesting insight into the different mechanisms underlying the DNA methylation-dependent gene expression of a particular tissue-specific gene across three mammalian species. Our study identified a conserved CpG island in the same region just upstream of the transcription start site in three mammalian SLC9C1 genes: mouse, rat, and human. Although this CpG island is maintained in all three genes, its specific role as a DNA regulatory element appears not to be completely conserved. Why and how this shift in DNA methylation-sensitive gene regulatory activity between the promoters and the upstream CpG islands occurred in mouse evolution is currently unclear, but this will be an interesting topic of future study. However, our study provides evidence that DNA methylation-dependent regulation of homologous genes can be maintained but the exact cis-regulatory elements responsible may not always be conserved across species.

## Acknowledgements

We would like to acknowledge funding support through the NIH 1 R15 HD101985-01A1. We would also like to acknowledge the Center for Bioinformatics and Functional Genomics (CBFG) at Miami University.

